# Differential Sequence Charge-clustering and Mixing-ratio Affect Stability and Dynamics of Heterotypic Peptide Condensates

**DOI:** 10.1101/2025.07.25.666616

**Authors:** Milan Kumar Hazra

## Abstract

Phase separation has emerged as a central mechanism cell utilizes to organize its material components and are extremely important in tuning various biological regulations. However, such condensates in cellular media is often very much heterogeneous in nature in terms of proteins and even polynucleotide’s presence in different concentrations. Present study explores stability and dynamics of heterotypic coacervates formed by polyampholyte binary peptides having differential charge-clustering limits. Systematic increase in differential charge-clustering in peptide pairs tunes the origin of heterogeneity and shows an enhanced stability of droplet than homogeneous ones with same average charge clustering of the two peptide pairs. In addition, stability of the condensate phase enhances linearly with the increase of high charge-clustering polymers in the system. Peptides with higher charge-clustering are 3-4 times slower diffusive within condensate phase than the lower charge-clustering ones due to heterogeneity in structural morphology of droplets, that too diminish as one lowers the difference of charge-clustering among sequence pairs forming the condensate. Coupled to the differential diffusivity of the polymers in condensates, droplet diffusion is nearly 7-35 times lower than bulk depending upon the mixing ratios of the polymers and variable sequence charge-clustering. Condensates with mild heterogeneity have shown an enhanced arrestation than the most heterogeneous ones originating from complementarity and better packing probed by energetics in the condensate. This study quantifies fundamental microscopic properties of heterotypic condensates formed through long-range electrostatic forces and particularly how they can be modulated by the differential charge-pattern in sequences and mixing fraction systematically.

## Introduction

Phase separation has emerged as a central mechanism through which cell accomplishes its compositional control spatio-temporally[1] and tunes various biological rhythms namely RNA-metabolism[2] and regulation, stress regulation[3], mechano-transduction[4], mitochondrial signaling[5], intra cellular storage[6] and even enhancement of concentration for substrates for specific biochemical reactions[7,8]. Even aberrant condensation leads to disease like states namely neurodegenerative ones[8,9] and even cancer[10]. It has been extensively shown that protein disorder plays a crucial role in recognition[11] and multi-body association of the assembly[12–15]. Given multicomponent nature and complex functionalities, these condensates are very much context dependent[16,17] as well as display distinct internal architecture[18,19]. Multivalent interactions[20–27] stemming from sequence organizational pattern[28,29] dictates such condensate’s stability and dynamics[29]. A sticker and spacer model successfully depicts organizational framework and sequence specific driving forces of phase separation[21,30–32]. Cross-linking of sticker motifs induce network mesh and binding energy of the peptides dictate the material properties and shape of the condensate[33,34]. Biomolecular condensates are complex fluids with extensive liquid-like nature spanning a broad range of visco-elasticity and are programmable as sequence features of the proteins involved alter degree and strength of cross-linking and leads to variations of viscoelastic nature of the network[35]. From an engineering viewpoint, condensates may be thought of tunable membrane-less organelles having significant soft colloid like character[36,37] and may be very much useful as an artificial carrier in intra-cellular cargo delivery, time dependent control and efficient release of cargo and even designing stimuli responsive artificial material[37]. And so, it is utmost important to decode the material properties as well as underlying physics of homo and heterotypic condensate formation involving multiple biologically relevant components namely proteins and polynucleotides as a function of the sequence characteristics[38,39].

Phase separation of various single component proteins with disordered domains has been studied in detail and points toward important role of various sequence features namely charge-clustering, sequence hydrophobicity, polar residues, cation-pi interactions in outlaying the formation kinetics, stability and dynamics of coacervates[27,28,30,32,40–42]. Several order parameters has been introduced in this regard namely sequence charge decoration[43], charge patten parameter[29] and fraction hydrophobicity[29] to delineate condensate’s features in a systematic way in analysis of intrinsically disordered protein’s conformational ensemble and connected efficiency toward homotypic phase separation. Composition as well as organization of amino acids in sequences affect stability and dynamics through a balance between enthalpy and entropic components in action[44,45]. As an example, elastin like hydrophobic protein domains induce entropy mediated phase separation as probed by direct NMR relaxation measurements and diffusion data[46]. Disorder plays an important role in formation of condensates as efficient and finite lifetime multivalent interactions are required for dynamic multi-body assembly[20]. This aspect is even connected to diffusivity inside the condensates and as has been shown Tau protein’s torsional fluctuation is inherently connected to liquid-like nature of assembly probed through picosecond time-resolved fluorescent depolarization measurements[47].

In the cellular context, condensates are often heterogeneous and not a single component phase and involve many components with diverse binding affinities to each other. Experimental studies with micro-rheology have shown significant sticker strength dependence of network reconfiguration timescale in ARG-GLY rich intrinsically disordered regions (IDRs) and single-stranded DNA in heterogenous condensates. Chain length of SS-DNA influences phase stability and internal viscosity but diffusivity of peptides is unaltered due to its variation[37]. Arrhenius law of activation energy has been observed to hold in predicting viscous nature of such heterotypic mesh[37]. Organizational heterogeneity has also been observed in heterotypic protein condensates of prion like domains in stress granule proteins namely FUS and hnRNPA. In 1:1 binary peptide system, heterotypic interaction has been reported to enhance the stability of condensate phase. The enhancement has been reported to be originating from complementary electrostatic interactions even in a system where only 10% of the residues are charged. α-synuclein and Prp forms a dynamic condensate with nanoscale electrostatic clusters within its microscopic assembly. Breaking and making of non-covalent interactions has been reported to influence the hierarchical structure formation of such condensates with diffusive nature[48].

Theoretical formulations and molecular simulations have extensively complemented the experimental understanding with all-atom or reduced resolution of representation of proteins and polynucleotides due to long-timescale required for such assembly’s equilibration[29– 32,42,49–56]. These studies however lack explicit effect of counter-ions or solvent and water’s role; they captured basic structural aspects as well as phase behaviours of relatively reduced miniature models of proteins and peptides. Sequence dependent re-entrant phase behaviour observed in patchy particles[57–59] and network-forming systems has also been observed in heterotypic protein-polynucleotide condensates and has shown important role in tuning dynamics and solubilization of droplets[60][61,62].

Present work sets the first step toward a systematic uncovering of sequence feature dependence of heterotypic phase separation. To that aim, we first plan to decode the effect of charge-clustering variability in sequences on stability and dynamics of phases formed by polyampholyte binary pair peptides. We offer a comparison of stability and internal dynamics, chain-reconfiguration timescales between homotypic and heterotypic phase separation of designed polyampholyte sequence pairs as a function of mixing ratios of peptides of interest as well as their charge-clustering variability to decode the effect of heterogeneity in condensate phase.

## Models and Simulations

### A. Designed Systems

To investigate how differential charge-clustering in sequence pairs and their mixing fractions alter stability and dynamics of binary protein coacervates we have designed three 40-residue sequence pairs that has been used in condensate formation with gradual enhancement of difference in charge pattern parameter (Δk) among the two sequences (Fig. 1). Representative condensates and their structural morphologies has been shown in Fig. 1 for Δk=0.86 and 0.30. While slate color denote the polymers with high-κ, limon green denotes low-κ ones. We have ensured that the average k of the two sequences is near 0.5 and compared our data with the condensates formed by homotypic phase separation of sequence κ=0.54. Several mixing fractions of the two polymers has been simulated namely *x*_*LOW−κ*_ = 0.10, 0.25,0.40,0.50,0.60,0.75 *and* 0.90. For clarity we are reporting only 5 of the simulated mixing fractions. We have ensured the neutrality of the sequences and the only order parameter that varies system-wide are sequence-specific charge-clustering and mixing fraction of the polymers.

**Figure 1:**
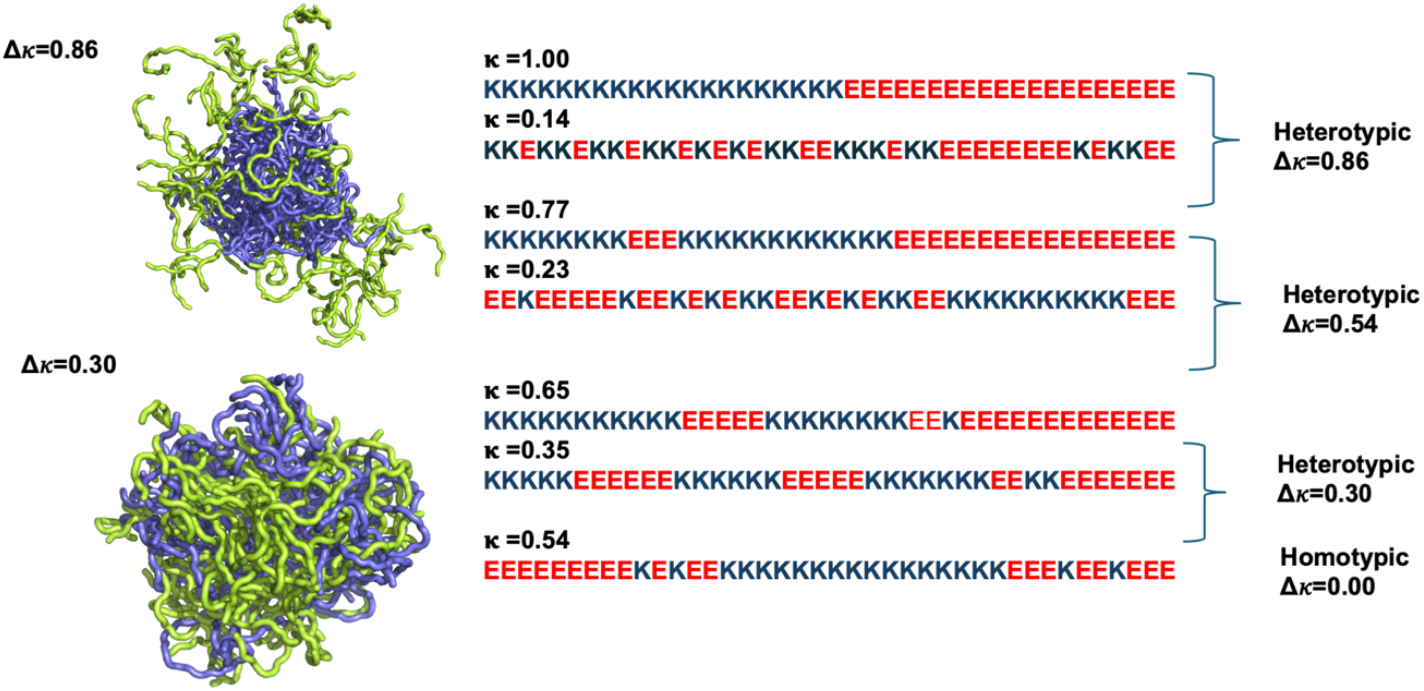
Designed sequence pairs with differential charge clustering. Heterotypic pairs with sequence κ=1.0 and 0.14 (Δκ=0.86), sequence κ=0.77 and 0.23 (Δκ=0.54) and sequence κ=0.65 and 0.23 (Δκ=0.30) has been shown along with a sequence that we have used to simulate homotypic condensate with κ=0.54 that matches the average of the heterotypic pairs. Representative condensates has been shown for Δκ=0.86 and Δκ=0.30 pairs at *x*_*LOW*κκ_ = 0.5 and T/Tc =0.4. Slate color represents polymers having high sequence κ and low-κ polymers has been shown by limon green.

### B. Simulation Details

As all-atom explicit solvent simulations are expensive to achieve timescales required to form mesoscopic condensates at equilibrium, a structure-based coarse-grained model[63][64,65] has been implemented to investigate equilibrated heterotypic condensates that allows comprehensive dissection of the effect of heterogeneity on condensate’s stability and dynamics. Each residue in the designed polyampholytes has been represented by a single bead and hasbeen assigned as either positively or negatively charged residue. The potential energy functional consist of the following terms, harmonic bonded and angular interactions to design the polymers, cosine dihedral interactions to impart stiffness of the polymers, electrostatic interactions among all charged beads, both intra- and inter-molecularly, and short-range dispersion interactions. We have not implemented any dihedral interactions as the designed peptides behave as IDRs.

An embedded implicit solvent model and salt effect has been used to screen electrostatic interactions between the charged beads through Debye-Hückel potential[64]. 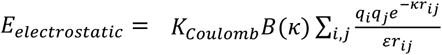; where *q*_*i*_ and *q*_*j*_ denote the charge of the *i*^th^ and j^th^ bead, *r*_*ij*_ denotes the inter-bead distance, *ε* is the solvent dielectric constant, and *K*_*Cloumb*_ = 4*π ε*_0_ =332 kcal/mol. *B*(*κ*) is a function of solvent salt concentration and the radius (*a*) of ions produced by the dissociation of the salt, an can be expressed as 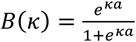. The Debye-Hückel electrostatic interactions of an ion pair act over a lengthscale of the order of *κ*^−1^, which is called the Debye screening length. K is related to ionic strength as,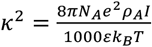; where *N*_A_ is the Avogadro number, *e* is the charge of an electron, *ρ*_A_ is the solvent density, *I* denotes solvent ionic strength, *k*_B_ is the Boltzmann constant, and T is the temperature. To avoid overlap among beads, a steep repulsion interaction has been defined as 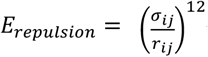 with *σ*_*ij*_=4 Å.

We have simulated 100 copies of binary polymer pairs at different mixing fractions as detailed above in a box of dimension 300 × 300 × 300 Angstrom^3^. Analysis of stability of the condensates and dynamics of polymers within the same has been quantified at ionic concentration 0.04 M with an implicit solvent medium of dielectric constant 80 and at multiple temperatures below critical point of the system. At each temperature, we have simulated multiple trajectories by solving Langevin equation to confirm convergence and averaging of the analysis shown here. A clustering algorithm has been used to identify condensates and the largest clusters from the trajectories.

## Results

### Sequence dependent phase stability in heterotypic condensates

To elucidate the effect of differential charge-clustering (as a measure of sequence heterogeneity) on droplet stability, temperature-density phase diagram has been plotted for three different sequence-pairs along gradual decrease in Δκ between two sequences in Fig 2 (A-C). Each panel shows five different mixing fractions of low and high-κ polymers and denoted with mixing fraction of the low-κ polymer *x*_*LOW*−*κ*_. At each timestep, we computed the density of polymers in condensate phase by counting the number of polymers beads in largest cluster within 5nm distance from center of mass of the cluster. We have employed a Cayley Tree like clustering algorithm that can capture the information of network mesh and can identify the largest cluster. The critical point has been determined by fitting density-temperature data to the Ising model with the following expression 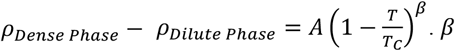 denotes critical exponent in Ising 3D model as 0.325. Stability of the condensate phase diminishes for the binary peptide κ=1.00 and 0.14 and κ=0.77 and 0.23 with the enrichment of low-k polymer in the system (Fig. 2A and B). Almost 3-4-fold decrease in condensate stability has been observed with increase in the fraction of low-κ polymer (here κ=0.14) from 0.10 to 0.90. Whereas moderate heterogeneity in charge-clustering of sequence-pairs (Fig. 1 panel C) enables tremendous stability of the condensate as one enhances fraction of low-κ polymers in condensate forming system at the expense of high-κ ones. We have observed only 1.5 times stability drop while mixing fraction of low-κ polymer reaches from 0.10 to 0.90 for Δκ=0.30 sequence pairs (Fig. 2C).

**Figure 2:**
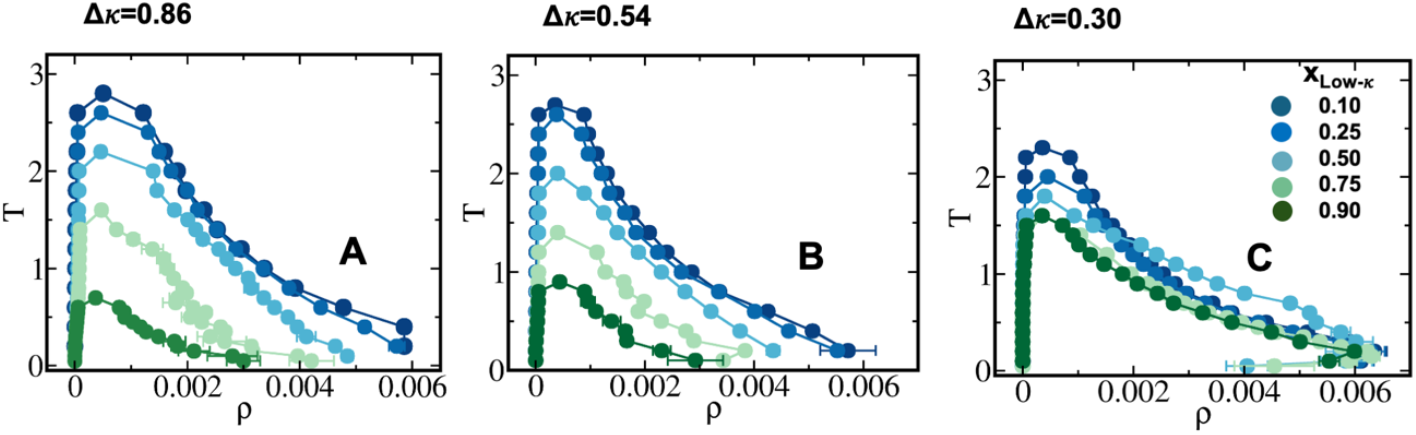
Phase diagram and criticality of the studied sequences. (A) Δκ=0.86 (sequence κ=1.0 and 0.14), (B) Δκ =0.54 (sequence κ=0.77 and 0.23) and (C) Δκ =0.30 (sequence κ=0.65 and 0.35). Different mixing fractions has been shown in each panel ranging from *x*_*Low*−*κ*_=0.1 to 0.9 with blue to green.

### Underlying energetics of phase separation and connection to origin of criticality

To decode origin of such heterogeneity induced stability-pattern in designed binary peptide condensates, total energy experienced by each polymer in the condensate phase has been computed as function of temperature scaled with respect to criticality and for all mixing fractions (Fig. 3). In addition, we have dissected the total energy into components namely interactions within high-κ polymers, low-κ polymer and cross interactions among polymers (Fig S1-S3). For highly heterogeneous condensates with sequence-pair κ=1.00 and κ=0.14, peptide-peptide interaction energy stabilizes the condensate phase up to 55 kcal/mol at a very low temperature far away from criticality (T/Tc=0.1) and that too decreases with mixing of low-κ polymers to 18 kcal/mol exactly 3-times as observed for phase diagram’s stability (Fig. 2A). Similarly, along temperature, the depletion of attractive energy is linear for extreme *x*_*LOW*−*κ*_ 0.1 and 0.9 due to presence of one-kind of polymer at its maximum but an enhanced stabilization has been observed in condensates formed by κ=1.00 and κ=0.14 sequence-pairs for mixing fraction *x*_*LOW*−*κ*_= 0.25 − 0.75 (where heterogeneity peaks in condensate phase) in the range T/Tc=0.2-0.6 due to temperature induced entropy in the system and a better mixing at relatively higher temperature at significant sequence heterogeneity. The stabilization loss along temperature also has a decreased gradient along mixing fraction ranging up to 3-fold electrostatic energy loss at *x*_*LOW*−*κ*_= 0.1 upon reaching critical point to 1.2-fold at extremely low-_*κ*_ polymer concentration in the droplet (*x*_*LOW*−*κs*_ 0.9). While with slightly reduced Δκ =0.54 induces stabilization up to 40kcal/mol for mixing fraction *x*_*LOW*−*κ*_= 0.1 at far away from criticality of the system (T/Tc=0.1), and nearly 18kcal/mol energetic stabilization has been observed for mixing fraction *x*_*LOW*−*κ*_= 0.9.

**Figure 3:**
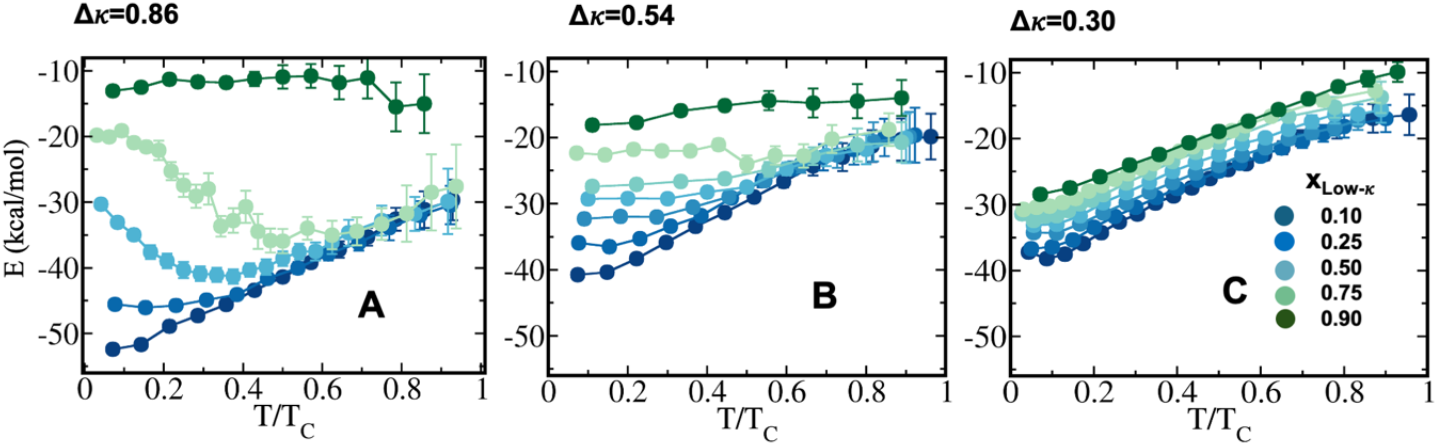
Energetic stabilization of heterotypic polymers in condensate phase. Average total energy, each polymer faces in condensate phase for sequence pairs with differential charge clustering (A) Δκ=0.86 (sequence κ=1.0 and 0.14), (B) Δκ=0.54 (sequence κ=0.77 and 0.23) and (C) Δκ=0.30 (sequence κ=0.65 and 0.35). Different mixing fractions has been shown in each panel ranging from *x*_*LOW*−*κ*_=0.1 to 0.9 with blue to green. Total energy a polymer experiences is composed of interactions faced from high-κ polymers and low-κ ones. Individual components has been shown in Fig. SI 1-3.

Whereas at mild heterogeneity of sequence-pairs (Δκ=0.30), an enhanced stabilization has been observed for the condensate systems with higher *x*_*LOW*−*κ*_ relative to Δκ =0.86 and 0.54 and the effect of mixing ratio (*x*_*LOW*−*κ*_) is low. The drop of energetic stabilization is nearly 1.3 times at T/Tc=0.1 compared to 3-times observed in Fig. 3A and 3B for condensates of sequence pairs with Δκ =0.86 and 0.54 along mixing fraction *x*_*LOW*−*κ*_ signifying a better complementarity among the sequence-pairs and effective packing that mixing ratios are unable to alter. This observation is in complete agreement with phase diagrams shown in Fig. 2. Along temperature, loss of energy in droplet phase is 2-3fold while reaching criticality (Fig. 3).

A dissection of pair-wise energies among the different kind of polymers has been shown in Fig S1-3 for Δκ =0.86, 0.54 and 0.30 respectively showcasing interactions among high-κ, low-κ and cross interactions among the two (Panel A, B and C respectively in each figure). Although the major gain in energy is mostly dominated by interactions among high-κ polymers at lower mixing ratio *x*_*LOW*−*κ*_ in all three Δκ systems (nearly 40kcal/mol at T/Tc=0.1) pointing to enhanced stability at *x*_*LOW*−*κ*_ as shown in Fig. 2, low-κ polymers contribute nearly 9-12 kcal/mol with its highest enrichment *x*_*LOW*−*κ*_ = 0.9 in the system (Fig. S1B-3B). In a way the energetic stabilization of the condensates stems from 10-18kcal/mol while the droplet is enriched with low-κ polymers and 40-50kcal/mol when enriched with high-κ polymers. Similar contribution has been observed from cross κ interaction energy at Δκ =0.86 and 0.54 at mixing fraction *x*_*LOW*−*κ*_ = 0.25 − 0.75 where mixing plays an important role. At Δκ=0.54 (Fig. S 3B), we have quantified a significant increase in interactions up to 25 kcal/mol among κ=0.35 polymers its maximum enrichment (*x*_*LOW*−*κ*_ = 0.9). In addition, interaction energy among κ=0.35 and κ=0.65 polymers also has increased during mixing ratio *x*_*LOW*−*κ*_ = 0.25 − 0.75 up to 15 kcal/mol signifying a better complementarity among the two sequences as pointed out earlier (Fig. S 3C).

### A comparative stability analysis of homotypic vs heterotypic phase separation

To decode the effect of heterotypic sequence pairs on phase stability, phase diagrams for all three Δκ sequence pairs has been shown along with a homogeneous condensate of sequence κ=0.54 (sequence has been shown in Fig. 1) at mixing fraction *x*_*LOW*−*κ*_ = 0.5. One must note a significant enhancement of stability has been observed for heterotypic sequence-pairs compared to homotypic one, while the homotypic condensate has critical point around reduced temperature 1.4, the heterotypic condensates have critical point 2-2.2 depending upon Δκ variation (Fig. 4A). We tried to map the correlation between occurrence of critical point and energy faced by each polymer in homotypic and heterotypic condensate phases along mixing fraction (Fig. 4B-C) at a fixed distance from criticality. The slope along *x*_*LOW*−*κ*_ gradually diminishes as one lower Δκ of sequence-pairs. One may also note a crossover of critical point of homotypic and heterotypic condensates at *x*_*LOW*−*κ*_ =0.80-0.90 regime when the stability of the dense phase is lesser than that of homotypic condensates. The same cross-over has also been observed in energy each polymer faces in droplet phase w.r.t mixing fraction *x*_*LOW*−*κ*_.

**Figure 4:**
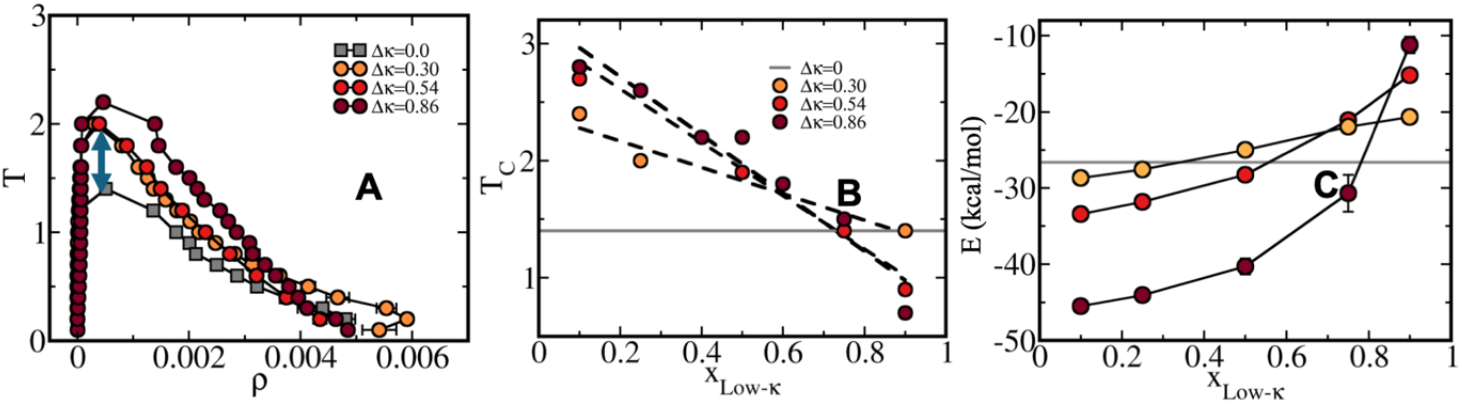
Comparative analysis of homotypic and heterotypic phase stability of condensates. (A) Comparative phase diagram for sequence pairs with Δκ=0.86 (Dark Red), Δκ=0.54 (Red) and Δκ=0.30 (Yellowish red) at *x*_*LOW*−*κ*_ =0.54 has been shown along with the same for homotypic κ=0.54 condensate (Grey squares). Enhanced stability has been observed for heterotypic ones and maximizes for highestΔκ. (B) Linear dependence of critical point along mixing fraction *x*_*LOW*−*κ*_ has been shown for condensate forming sequence pairs Δκ=0.86 (Dark Red), Δκ=0.54 (Red) and Δκ=0.30 (Yellowish red) and compared with homogeneous κ=0.54 condensate (Grey line). (C) Average total energy experienced by polymers in condensate phase at constant T/Tc =0.4 along mixing ratio *x*_*LOW*−*κ*_ for various Δκ variant pairs.

### Liquid-like nature and translational diffusivity in condensate phase

To quantify the internal dynamics within heterotypic condensates, we measure the diffusion coefficients of high and low-κ polymers while within the droplet at different temperatures approaching criticality. The diffusion coefficients have been evaluated as the slope of mean squared displacement (Fig. S4 for MSD of polymers in droplet phase) computed as 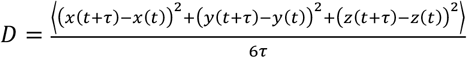. While calculating the diffusion coefficients of polymers, we have ensured that the polymers remain in the droplet during the entire time the mean squared displacement (MSD) is being computed. Fig. 5 (A-C) plots the MSD of polymers in droplet phase for Δκ=0.86, 0.54 and 0.30 respectively at T/Tc =0.4 and *x*_*LOW*−*κ*_ = 0.50. While solid line represents the MSD of high-*κ* polymers, dotted curve represents the same for low-*κ* ones. It is interesting to note as Δκ of the sequence pairs decrease the two MSDs approach each other and mediate a dynamic homogeneity for Δκ=0.30. Representative trajectories has been shown for low-κ (green) and high-κ polymers (red) in condensate phase among all the polymers’ trajectories (Grey) at T/Tc=0.2. We observe low-κ polymers form the surface of droplet and move on the surface with significantly higher diffusivity when heterogeneity in sequence pairs is high enough while high-κ ones form the core. With diminishment ofΔκ, a gradual penetration of low- κ polymers in the droplet is visible even from trajectory. Fig. 6 (A-C) plots the ratio of diffusivity for low-κ and high-κ polymers 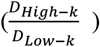 along temperature scaled to critical point for all the mixing fractions. For Δκ=0.86, low-*κ* polymers diffuse in condensates almost 4-fold higher than high- *κ* ones at far away from criticality (T/Tc=0.1) due to an enhanced interaction of high-*κ* polymers at lower mixing fraction (*x*_*LOW*−*κ*_) and this ratio diminishes gradually to 1 as temperature approaches critical point. For Δκ=0.54 sequence pairs in the condensate phase, the same ratio tends to be maximum at T/Tc=0.1 being 2-3 times than that of high-κ polymers subject to mixing ratio. As the mixing ratio *x*_*LOW*−*κ*_ enhances, the slope of the ratio with respect to T/Tc gradually decreases. At a given temperature scaled by Tc, 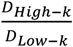 for Δκ=0.86 and 0.54 has higher values as *x*_*LOW*−*κ*_ tends to 0 enriching the droplet with high-κ polymers. Surprisingly at moderate heterogeneity of sequence pairs Δκ=0.30, we have observed both low-κ and high-κ polymers have nearly similar translational diffusivity irrespective of mixing ratio and temperature. A better packing and enhanced cross κ polymer interactions drive such scenario.

**Figure 5:**
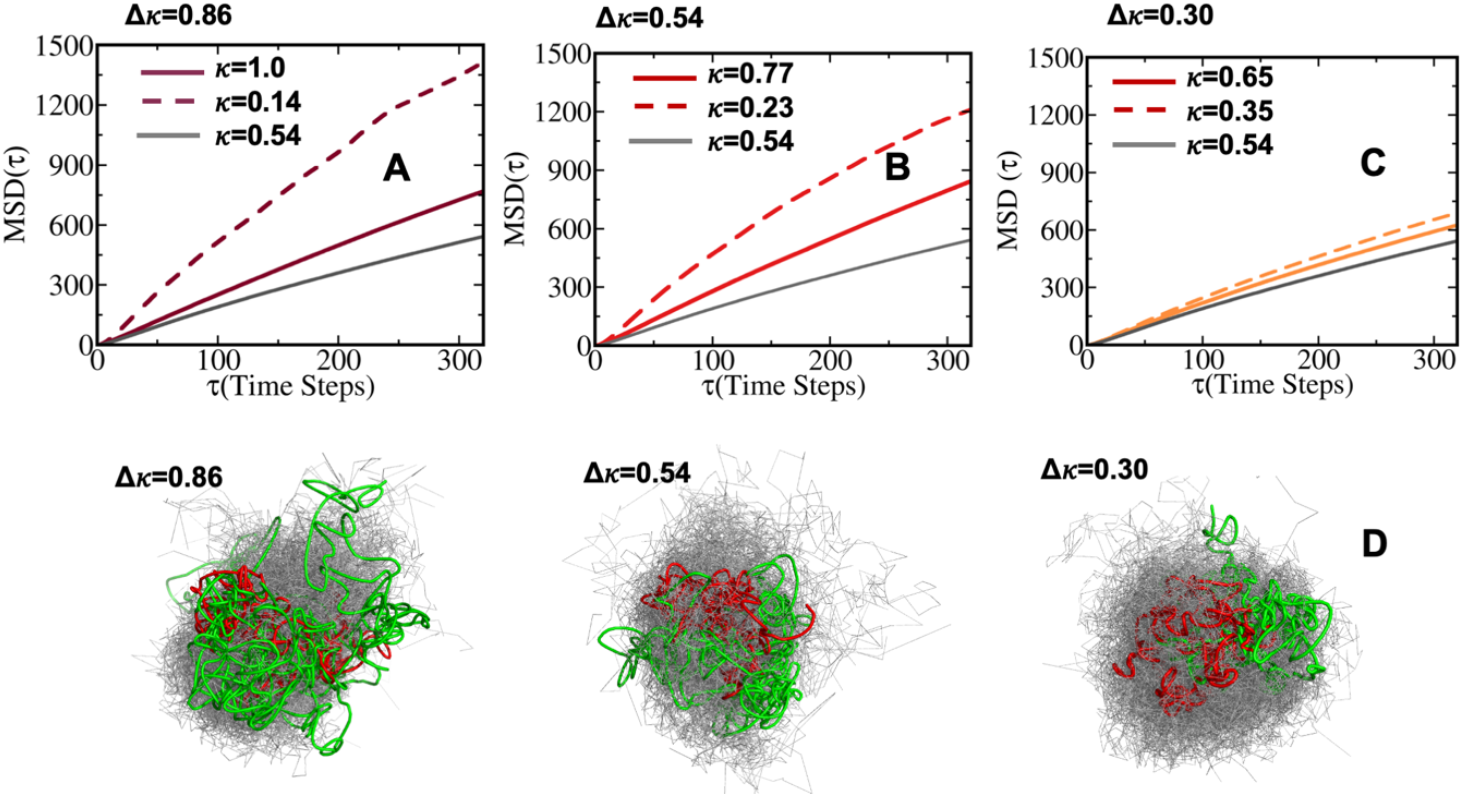
Translational diffusivity in condensates and effect of heterogeneity. Mean squared displacement (MSD) of polymers has been shown for sequence pairs with (A) Δκ=0.86 (Dark Red), (B) Δκ=0.54 (Red) and (C) Δκ=0.30 (Yellowish red) at mixing ratio *x*_*LOW*−*κ*_= 0.5 and at T/Tc=0.4. While solid line indicates MSD for polymers with high charge clustering, dashed line indicates MSD for polymers with lower charge clustering. Grey curve indicates the same for homogeneous condensate with average κ=0.54. Representative trajectories in condensate phase for high-κ polymer has been shown with red and for low-κ polymer with gsreen in all polymer’s trajectories (grey) in condensates formed by Δκ=0.86, 0.54 and 0.30 respectively.

**Figure 6:**
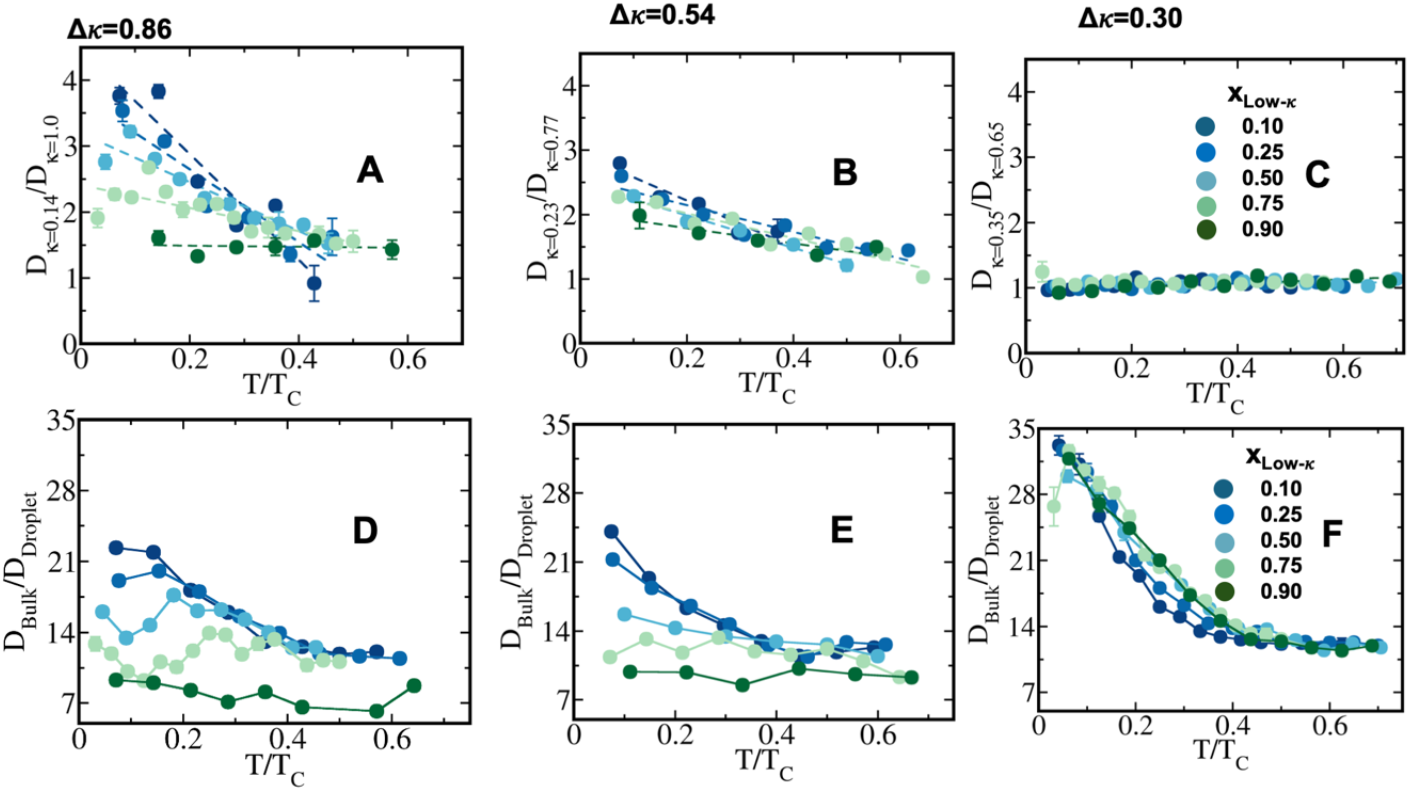
Differential dynamics in condensate phase and bulk. Ratio of diffusivity between low-κ and high-κ 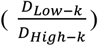 polymers in condensate phase along temperature scaled to criticality for sequence pairs (A) Δκ=0.86 (sequence κ=1.0 and 0.14), (B) Δκ=0.54 (sequence κ=0.77 and 0.23) and (C) Δκ=0.30 (sequence κ=0.65 and 0.35). Polymer diffusivity at dense phase compared to bulk 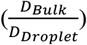 along temperature scaled to criticality of droplet phase for sequence pairs (D) Δκ=0.86 (sequence κ=1.0 and 0.14), (E) Δκ=0.54 (sequence κ=0.77 and 0.23) and (F) Δκ=0.30 (sequence κ=0.65 and 0.35). Different mixing fractions has been shown in each panel ranging from *x*_*LOW*−*κ*_ =0.1 to 0.9 with blue to green.

Average diffusivity in bulk is nearly 7-20 times higher than condensate phase subject to different mixing fractions and temperature for Δκ=0.86 and 0.54 (Fig. 6 panel D and E). However, the diffusion is liquid-like for condensates formed by all three Δκ pairs. For condensates formed by Δκ =0.86 and 0.54 sequence pairs,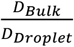 gradually decrease along *x*_*LOW*−*κ*_ indicating enhanced mobility in the dense phase correlated to lower stability (Fig. 2A and B) of the same. However, for condensates formed by sequence pairs κ=0.30, we have observed an extensive dropdown of diffusivity in the condensate phase and due to which bulk diffusivity is nearly 14-35 times higher than droplet phase subject to temperature variation (Fig. 6 panel F). We have observed no evidence of the effect of mixing on 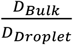 as well as in previous analysis.

### Reconfiguration Lifetime in Dense Phase

To decode the relation between average translational diffusivity to single-molecule conformational kinetics of polymers in condensate, we have computed polymer end-to-end distance vector’s time correlation function as follows: 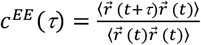 where 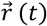 represents end-to-end distance vector of peptides (Fig. S 5A-C). A bi-exponential function has been fitted to get the average time-constants for end-to-end distance dynamics of polymers. Average timescale of correlated end-to-end distance fluctuation in condensate phase for high-κ and low-κ polymers has been shown in Fig. 7 (panels A-C) as 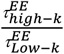. We have observed a 1.5-12-fold enhanced slowdown of end-to-end distance vector dynamics for high-κ ones with respect to low-κ ones subject to increased heterogeneity (Δκ), temperature and mixing fractions. For condensate formed by sequence pairs Δκ=0.86, we find 10-times slowdown for high-κ polymers with respect to low-κ one at T/Tc=0.1 for profound population of high-κ polymers (Fig. 7A). A slowdown of nearly 6-fold has been observed at similar conditions for Δκ=0.54 (Fig. 7B). As temperature approaches criticality the ratio tends to decrease and around T/Tc=0.6 we observe the slowdown for high-κ one is nearly 1.5 to 3-times that of low-k polymers for both Δκ=0.86 and 0.54 pairs. As low-κpolymer enriches the droplet we have a gradual decrease in the ratio at same distance from critical point. For mixing ratio 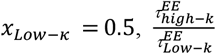 is nearly 6 at T/Tc=0.1 and drops to 3 at T/Tc=0.6 indicating enhanced dynamics. 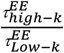behaves like 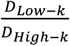 along temperature scaled to Tc but while translational diffusivity obtains a maximum 4-times slowdown for high-κ ones with respect to low-κ polymers, single-molecule dynamics reaches 9-10 times slowdown at far away from critical point. While for Δκ=0.30, as observed earlier, in translational diffusion, we have observed similar chain reconfiguration timescale for both low and high-κ polymers irrespective of temperature and mixing in accordance to Fig. 6C of translational diffusivity.

**Figure 7:**
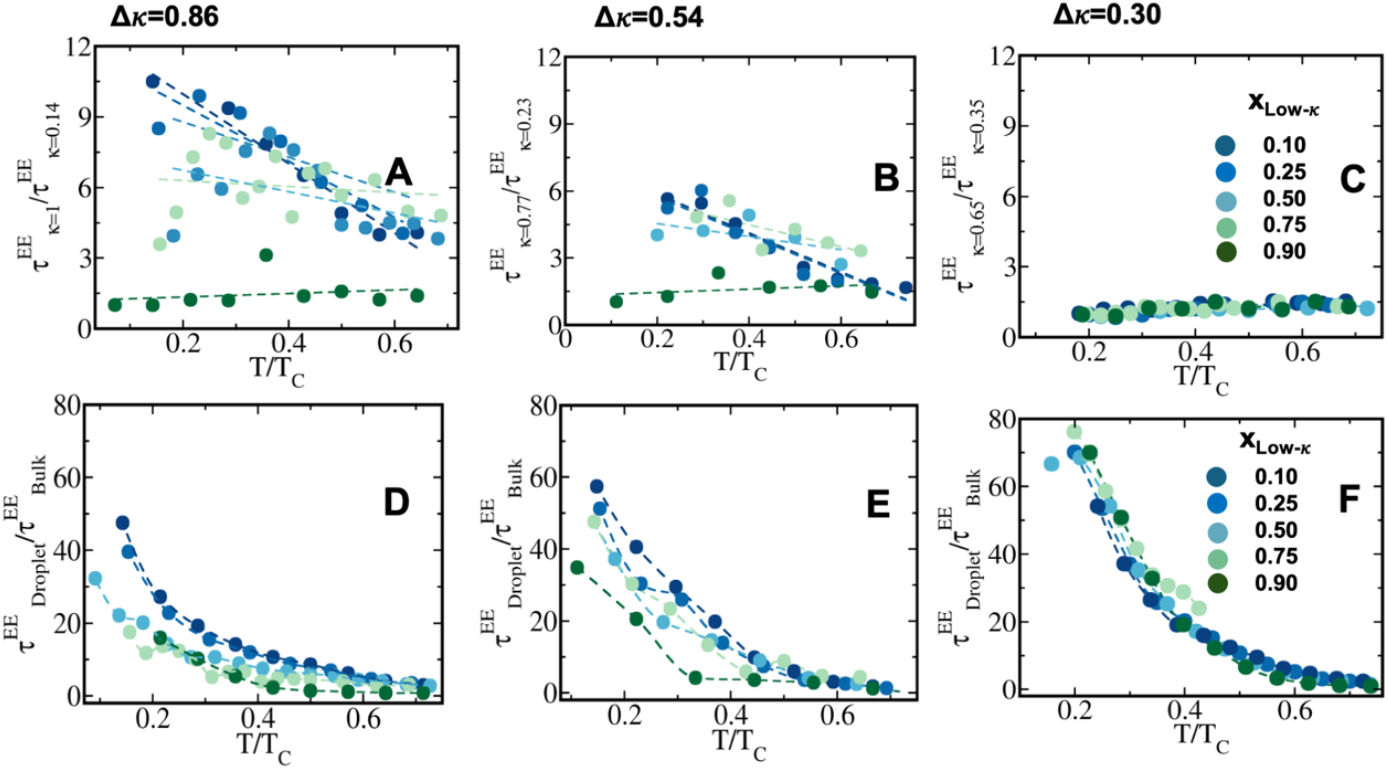
Chain reconfiguration dynamics in condensates. Average chain reconfiguration lifetime compared among low-κ and high-κ 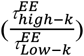 polymers in condensate phase along temperature scaled to criticality for sequence pairs (A) Δκ=0.86 (sequence κ=1.0 and 0.14), (B) Δκ=0.54 (sequence κ=0.77 and 0.23) and (C) Δκ=0.30 (sequence κ=0.65 and 0.35). Chain reconfiguration timescale of polymers compared to bulk 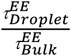 along temperature scaled to critical temperature for droplets of sequence pairs (D) Δκ=0.86 (sequence κ=1.0 and 0.14), (E) Δκ=0.54 (sequence κ=0.77 and 0.23) and (F) Δκ=0.30 (sequence κ=0.65 and 0.35). Different mixing fractions has been shown in each panel ranging from *x*_*LOW*−*κ*_ =0.1 to 0.9 with blue to green.

End-to-end distance vector dynamics in droplet phase is 20-60-fold slower than bulk for Δκ=0.86 and the slowdown is nearly 40-60 times for Δκ=0.54 subject to various mixing fractions at farthest away from criticality (Fig. 7 panel D and E and Fig. S 5D-E). As temperature tends to move toward criticality 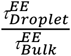 drops to 1. Like previous analysis of translational diffusivity in condensate phase of sequence pairs with mild heterogeneity Δκ=0.30, we observed an enhancement of slowdown up to 70-80 times that of bulk for end-to-end distance vector reconfiguration at far away from criticality T/Tc=0.1 and again no effect of mixing fraction has been observed (Fig. 7 panel F and Fig. S 5F).

To elucidate the effect of heterogeneity of sequence pairs and mixing fraction on single-molecule dynamics, translational diffusivity and to compare the same with homogeneous condensate’s dynamics, 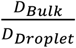 as a function of mixing fraction *x*_*LOW*−*κ*_ has been shown in Fig.

8A and 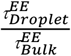 in Fig. 8B for constant T/Tc=0.2. Grey line depicts the same for homogeneous condensates of sequence κ=0.54 with Δκ=0.00. 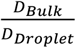 and 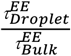 shows a steady decrease along enhancement of *x*_*LOW*−*κ*_ for Δκ=0.86 and 0.54 while Δκ=0.30 shows no dependence on mixing. Ratio of bulk to droplet diffusivity decrease from 20-fold to 10-fold while one moves from *x*_*LOW*−*κ*_= 0.1 to 0.9 at constant T/Tc= 0.2 showing enhanced mobility along enrichment of low-κ polymers and correlated to critical point’s variation shown in Fig. 2B showing a linear connection to stability of dense phase. Whereas 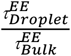 decreases from 40-fold to15-fold as one enriches the droplet with low-κ polymers for significantly high heterogeneity in sequence charge-clustering of the pairs.

**Figure 8:**
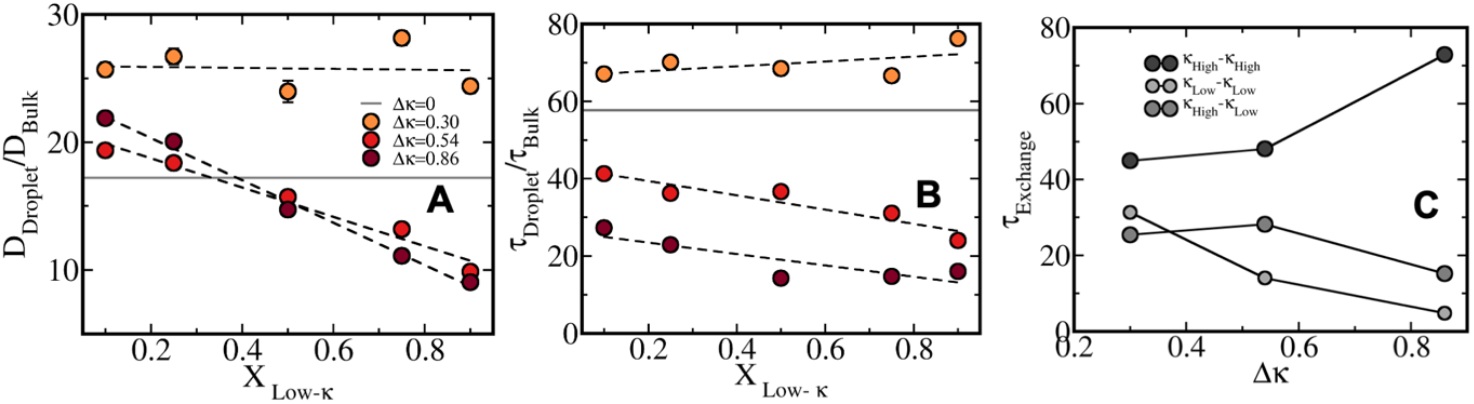
Heterogeneous dynamics in condensate along mixing fraction. (A) Diffusivity ratio in droplet phase and bulk for polymers and (B) chain reconfiguration timescale of polymers in droplet in bulk phase for sequence pairs Δκ=0.86 (sequence κ=1.0 and 0.14, Dark Red), Δκ=0.54 (sequence κ=0.77 and 0.23, Red) and Δκ=0.30 (sequence κ=0.65 and 0.35, Yellowish Red) along mixing fraction *x*_*LOW*−*κ*_ at T/Tc=0.4. Grey line indicates the same for homogeneous condensates with sequence κ=0.54. (C) Average lifetime of interchain contact among high-κ polymers, low-κ ones and cross interactions at absolute temperature T*=0.4 for mixing fraction *x*_*LOW*−*κ*_ = 0.5 along sequence heterogeneity Δκ.

Single polymer conformational kinetics in droplet phase as well as its translational diffusion are inherently connected to average lifetime of inter-chain contacts in the droplet phase among polymers (Fig. S6). To elucidate the timescale through which a polymer stays in contact with another polymer in droplet phase, we introduce a time correlation function 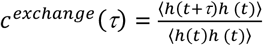 ; h(t) denotes a step function; h(t)=1 when two polymers are neighboring ones for a time frame and 0 otherwise (Fig. S6 A-C). We show the variation of average timescale of polymers in contact (τ_*Exchange*_) among high-κ polymers, low-κ and cross-κ ones for a specific temperature T*=0.4 and at mixing fraction 0.5 along heterogeneity of the sequence pairs (Δκ) inducing condensation. We have observed an increase in τ _*Exchange*_ for high-κ polymers in contact as one increase heterogeneity of charge-clustering in condensates. With increase in heterogeneity the sequence κ also enhances for relatively high-κ polymers. While τ_*Exchange*_ for the low-κ one behaves exactly opposite to that of high-κ one along Δκ as the sequence k for these low-κ ones move toward lower values and leading to easily breakable contacts. While timescale of contact among cross-κ polymers at lower Δκ regime follows the high-κ behavior and as Δκ increases follows the trend of low-κ ones.

### Droplet Shape Anisotropy

To correlate the stability pattern of condensate phase to the geometry of the same, we estimate shape anisotropy of condensates from a sphere. Shape anisotropy has been defined as 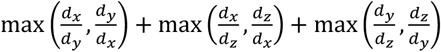 where *d*_*x*_, *d*_*y*_, *d*_*z*_ are the largest diameter in each dimension of the largest cluster. Along this definition, an ideal sphere has the value of 3. Deviation from the same indicates a distorted condensate from sphere. Fig. 9A plots the relative anisotropy from a spherical condensate along mixing ratios at T/Tc=0.4 for Δκ=0.86, 0.54 and 0.30 respectively. At lower mixing fraction, all Δκ variants have relatively similar shape anisotropy and enhances as *x*_*LOW*−*κ*_ increases for Δκ=0.86, 0.54 and remains nearly similar for Δκ=0.30 denoting complementarity of sequence pairs in similarity to energy analysis in Fig. 3 and Fig. S3. One must note the complementarity is maximum for homogeneous condensates and has minimum anisotropy in the studied systems (shown in Grey). Along temperature the anisotropy is increasing as the stability of the dense phase diminish for heterotypic ones for mixing fraction *x*_*LOW*−*κ*_ =0.5 and homotypic one shows fair stability within the time window of T/Tc =0.2 to 0.7 (Fig. 9B and Fig. S7).

**Figure 9:**
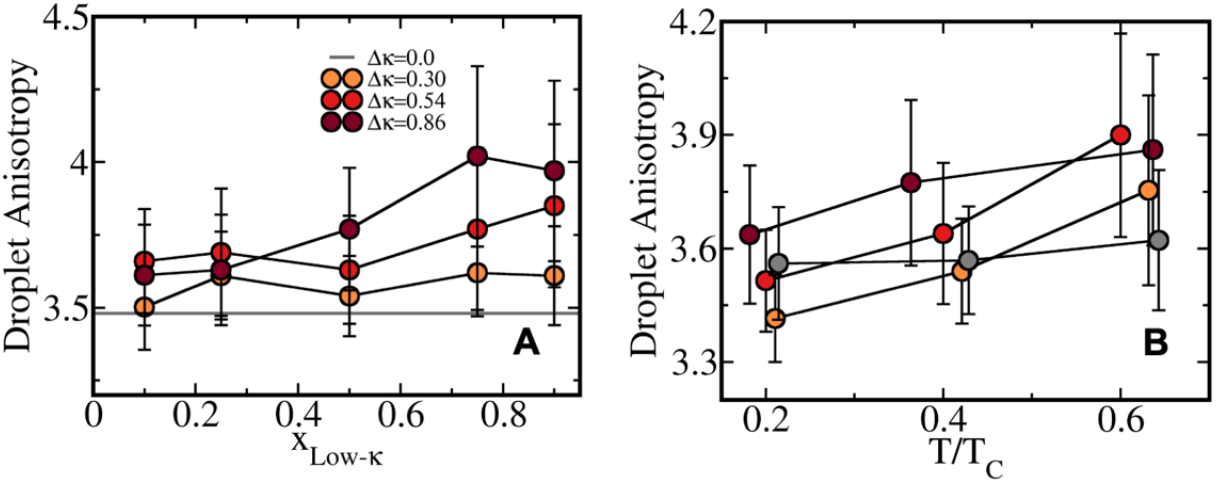
Anisotropy in geometry of condensate. **(A)** Shape anisotropy parameter shown along *x*_*LOW*−*κ*_ at fixed T/Tc=0.4. Grey line indicates the same for homogeneous condensates with sequence κ=0.54 for comparison. (B) Shape anisotropy parameter along scaled temperature to criticality in the range 0.2-0.7 for condensates formed by sequence pairs Δκ=0.86 (sequence κ=1.0 and 0.14, Dark Red), Δκ=0.54 (sequence κ=0.77 and 0.23, Red) and Δκ=0.30 (sequence κ=0.65 and 0.35, Yellowish Red).

Visual representation of structural morphology and heterogeneity along with droplet shape has been shown in Fig. 10 for two extreme mixing fractions *x*_*LOW*−*κ*_ =0.25 and 0.75 and for extremely heterotypic Δκ=0.86 (Fig. 10 A and B) and mild heterotypic Δκ=0.30 sequence-pairs (Fig. 10 C-D). It is very interesting to note that at higher heterogeneity of sequence pairs Δκ=0.86, the low-κ polymer forms the surface of the dense phase while the high-κ one forms the core. The differential diffusivity obtained in Fig. 6(A-C) and end-to-end distance vector dynamics Fig. 7 (A-C) is essentially related to the structural morphology and surface diffusivity.

**Figure 10:**
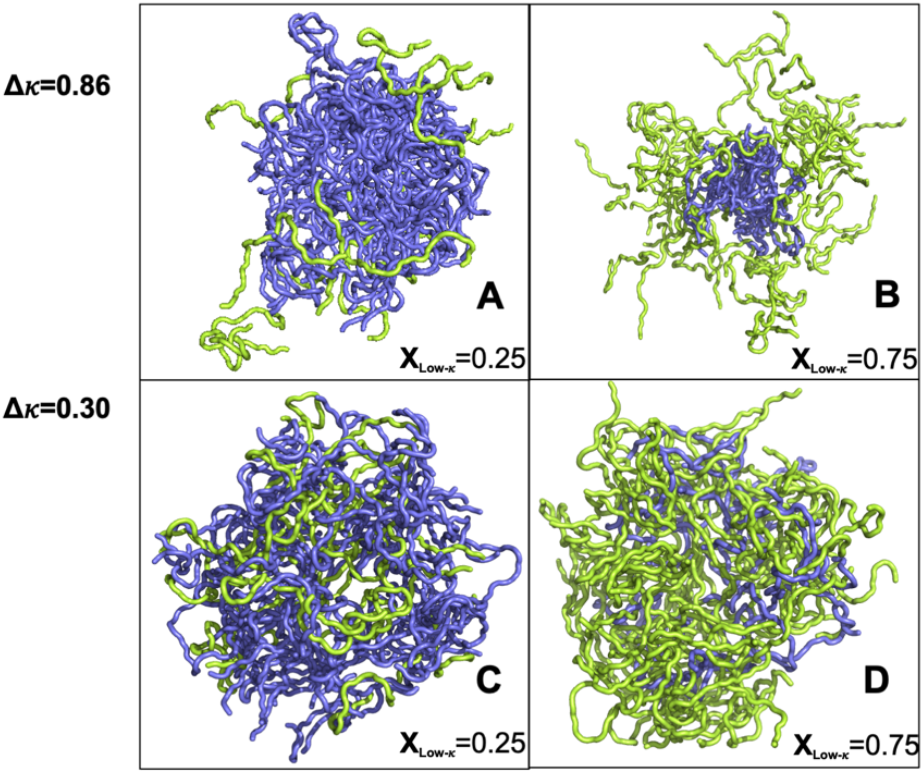
Representative snapshots of condensates delineating structural morphology and related geometric features for (A) lower population of low-κ polymer for highly heterotypic sequence pairs (Δκ=0.86), (B) higher population of low-κ polymer for highly heterotypic sequence pairs (C) lower population of low-κ polymer for mild heterotypic sequence pairs (Δκ=0.30), (D) higher population of low-κ polymer for mild heterotypic sequence pairs.

As the heterogeneity in pairs of sequence lowers, low-κ polymer penetrates the droplet and lowers the dynamic heterogeneity shown in Fig. 10 (C-D)

## Conclusions

In conclusion, present work lays the foundation for a systematic investigation into how sequence features influence heterotypic protein phase separation. Toward our initial goal to unravel how variations in charge-clustering within binary sequences affect stability and dynamics of phases, we have showcased temperature-density phase diagram, translational dynamics in condensates, and chain reconfiguration timescales of homotypic versus heterotypic phase separation in designed polyampholyte sequence pairs, examining how these properties vary with peptide mixing ratios and charge-clustering differences. A detailed microscopic view of relative composition, sequence heterogeneity driven altered structural morphology of condensate phase and fascinating connection of stability and dynamics profile with altered condensate morphology has been elucidated. Among many valuable outcomes, we have observed enhanced stability of condensates with significantly different charge clustering in binary peptides with respect to its homotypic counterpart having one kind of sequence with charge-pattern parameter similar to the same for binary peptides involved in heterotypic condensates. However, better packing and interaction energy in mild heterotypic condensates favor an enhancement of droplet stability when it has enriched population of low-κ polymers which is not the case for higher charge-clustering heterogeneity in binary peptides. Differential charge-clustering influences structural morphology of the formed condensate and enhances shape anisotropy rapidly with temperature. At relatively higher difference in charge-clustering of the two sequences forming condensate, one finds high-κ peptide forms the core of the droplet while low-κ ones are forming surface leading toward a 3-4 times higher diffusivity of low-κ peptides over high-κ ones due to surface rolling. As the charge-clustering heterogeneity diminish, the low-κ polymers penetrates more into the droplet phase and nearly has similar diffusivity to high-κ ones. Average timescale of chain reconfiguration is nearly 3-12 times faster for low-κ polymers being on the surface than high-κ ones forming core and subject to extent of composition and sequence charge-clustering heterogeneity. Overall diffusivity in condensate phase is 7-35 times slower with respect to bulk is in accordance with a slowdown of 5-70 times in chain reconfiguration timescale at far away from criticality and subject to variation of differential charge clustering and their mixing fraction. In short, our study delineates the effect of charge-clustering heterogeneity on binary peptide’s condensate forming ability and its inherent dynamics leading to the first step of systematic understanding of heterotypic condensation. Short-range hydrophobic and cation-ρε interactions and their different extent in sequence pairs will be examined in the upcoming study to elucidate the interplay of long- and short-range interactions in heterotypic mesh.

## CRediT authorship contribution statement

Milan Kumar Hazra: Conceptualization, Methodology, Investigation, Formal analysis, Writing – original draft, Review, Editing, Acquiring Funding.

## Acknowledgement

This work was funded by seed grant from IIT Jodhpur (I/SEED/MKZ/20230169). MKH also thanks High Performance Computing facility at IIT Jodhpur for computational support.

