## Supplemental Figure for "Differential Sequence Charge-clustering and Mixing-ratio Affect Stability and Dynamics of Heterotypic Peptide Condensates"

Milan Kumar Hazra*

Department of Chemistry

Indian Institute of Technology, Jodhpur

NH 62, Surpura Bypass Rd, Karwar, Jheepasani, Rajasthan

INDIA, 342030

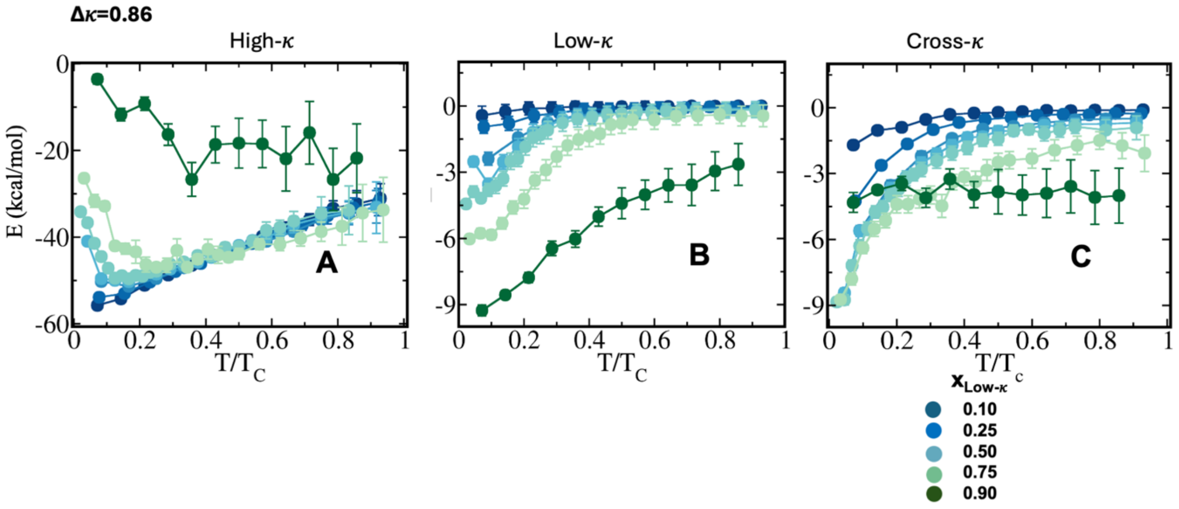


**Figure S1: Dissection of total energy in self- and cross- interaction energy components in heterotypic protein condensates of** sequence pairs κ=1.0 and 0.14**.** (A) Electrostatic energy among high-κ peptides that each polymer faces in condensate phase (B) Average electrostatic energy among Low-κ peptides that each polymer faces in condensate phase (C) Average electrostatic cross interaction energy among low and high-κ peptides that each polymer faces in condensate phase for sequence pairs with differential charge clustering Δκ=0.86 (sequence κ=1.0 and 0.14). Different mixing fractions has been shown in each panel ranging from $x_{Low-\kappa}$=0.1 to 0.9 with blue to green.

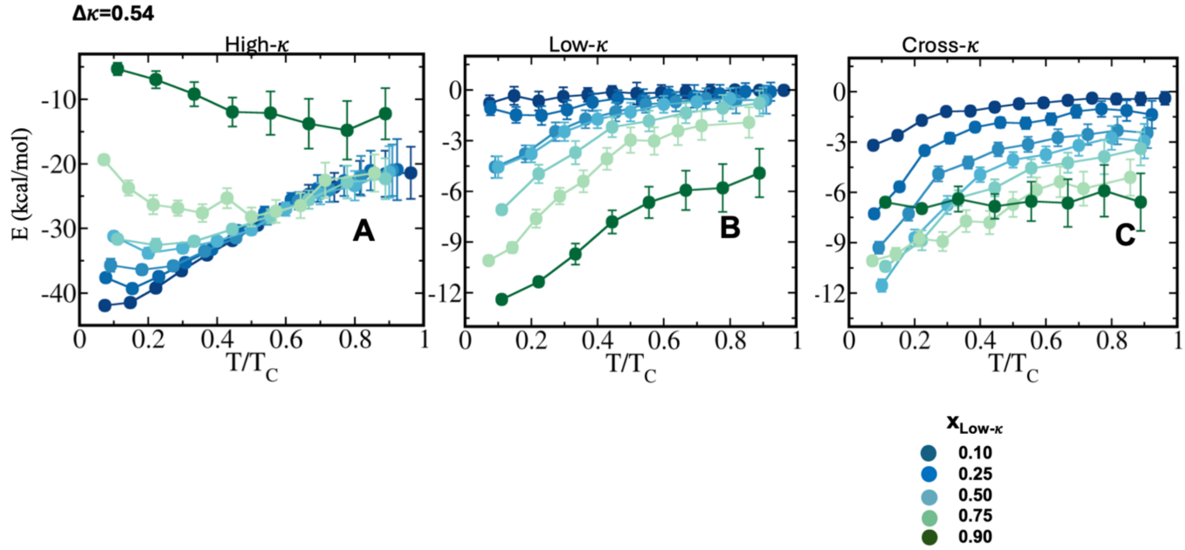


**Figure S2: Dissection of total energy in self- and cross- interaction energy components in heterotypic protein condensate of** sequence pairs κ=0.77 and 0.23**.** (A) Electrostatic energy among high-κ peptides that each polymer faces in condensate phase (B) Average electrostatic energy among Low-κ peptides that each polymer faces in condensate phase (C) Average electrostatic cross interaction energy among low and high-κ peptides that each polymer faces in condensate phase for sequence pairs with differential charge clustering Δκ=0.54 (sequence κ=0.77 and 0.23). Different mixing fractions has been shown in each panel ranging from $x_{Low-\kappa}$=0.1 to 0.9 with blue to green.

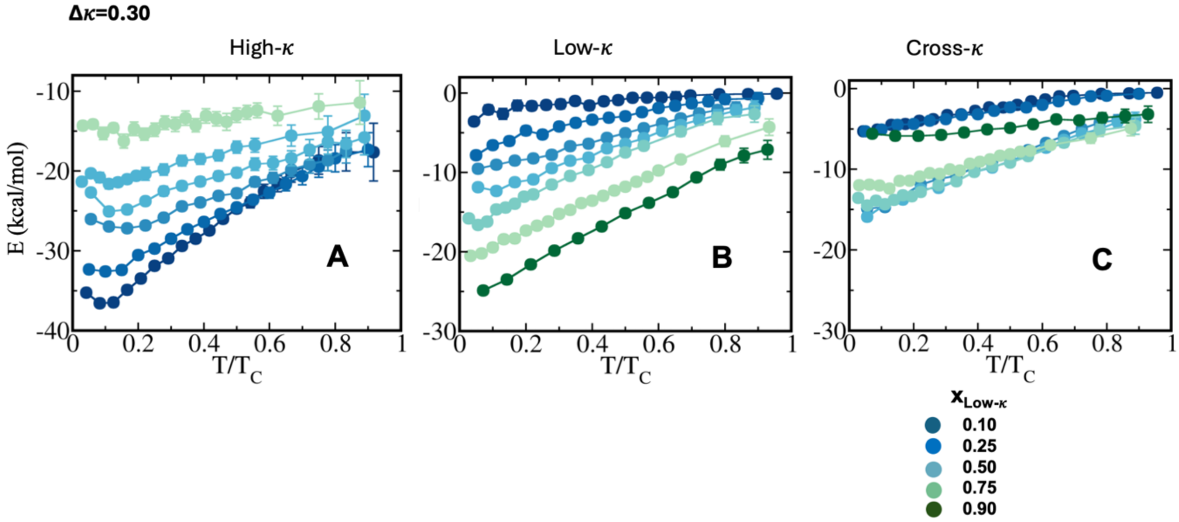


**Figure S3: Dissection of total energy in self- and cross- interaction energy components in heterotypic protein condensate of** sequence pairs κ=0.65 and 0.35**.** (A) Electrostatic energy among high-κ peptides that each polymer faces in condensate phase (B) Average electrostatic energy among Low-κ peptides that each polymer faces in condensate phase (C) Average electrostatic cross interaction energy among low and high-κ peptides that each polymer faces in condensate phase for sequence pairs with differential charge clustering Δκ=0.54 (sequence κ=0.65 and 0.35). Different mixing fractions has been shown in each panel ranging from $x_{Low-\kappa}$=0.1 to 0.9 with blue to green.

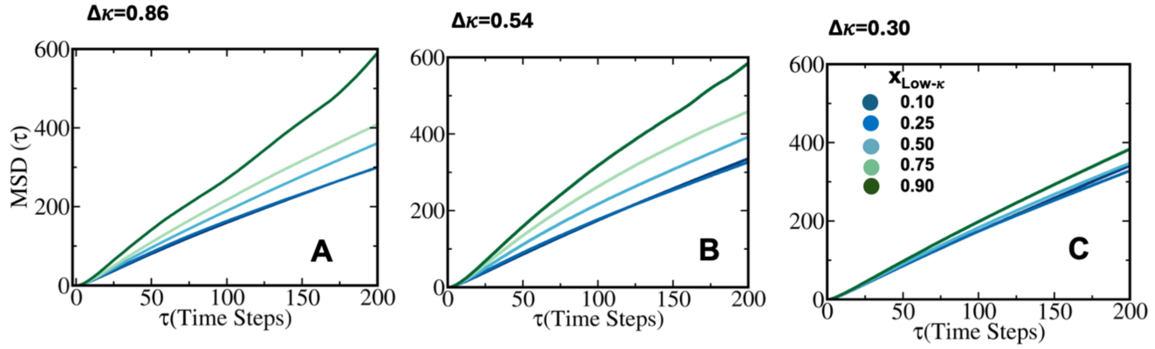


**Figure S4: Average mean squared displacement (MSD) of polymers in droplet phase for differential sequence pairs at absolute T*=0.6 (A)** Δκ=0.86 (sequence κ=1.0 and 0.14) (B) Δκ=0.54 (sequence κ=0.77 and 0.23) (C) Δκ=0.30 (sequence κ=0.65 and 0.35). Different mixing fractions has been shown in each panel ranging from $x_{Low-\kappa}$=0.1 to 0.9 with blue to green.

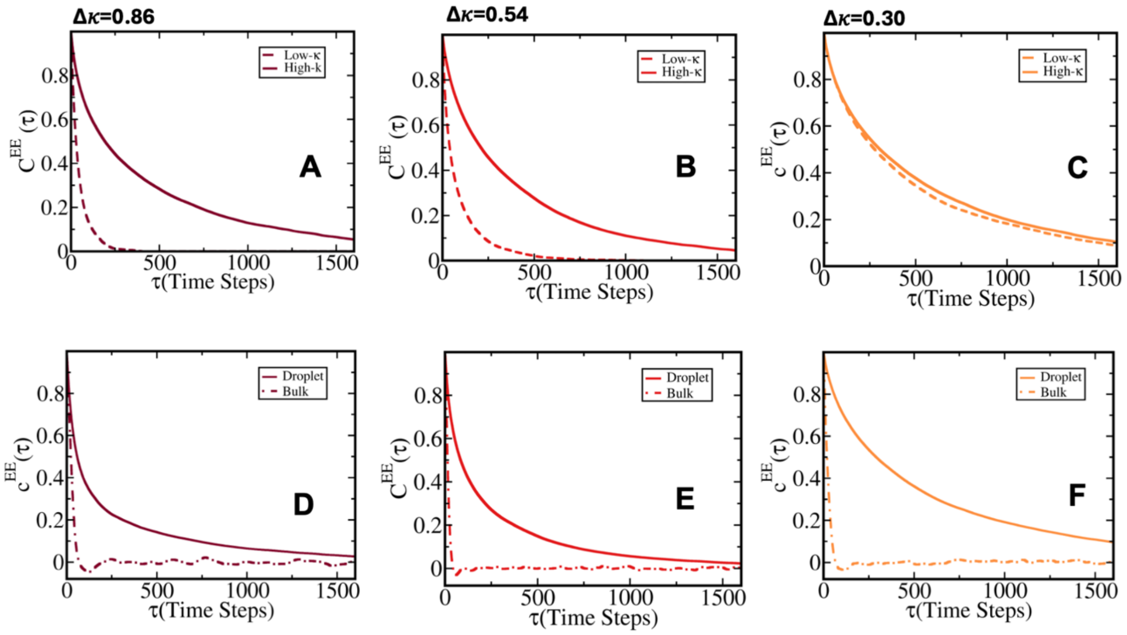


**Figure S5: Chain reconfiguration time correlation analysis in droplet and bulk phase.** Representative time correlation of end-to-end distance vector for low-κ and high-κ polymers in condensates at T/Tc=0.4 and mixing ratio $x_{Low-\kappa} = 0.5$ with sequence pairs (A) Δκ=0.86 (sequence κ=1.0 and 0.14) (B) Δκ=0.54 (sequence κ=0.77 and 0.23) (C) Δκ=0.30 (sequence κ=0.65 and 0.35). While solid line represents correlation for high-κ polymers, dashed line represent low-κ polymers. Representative average time correlation of end-to-end distance vector for polymers in condensate and bulk at T/Tc=0.4 and mixing ratio $x_{Low-\kappa} = 0.5$ with sequence pairs (D) Δκ=0.86 (sequence κ=1.0 and 0.14) (E) Δκ=0.54 (sequence κ=0.77 and 0.23) (F) Δκ=0.30 (sequence κ=0.65 and 0.35). While solid line represents correlation in droplet polymers while dot-dashed line represent the same in bulk. Color scheme represent individual Δκ variants respectively.

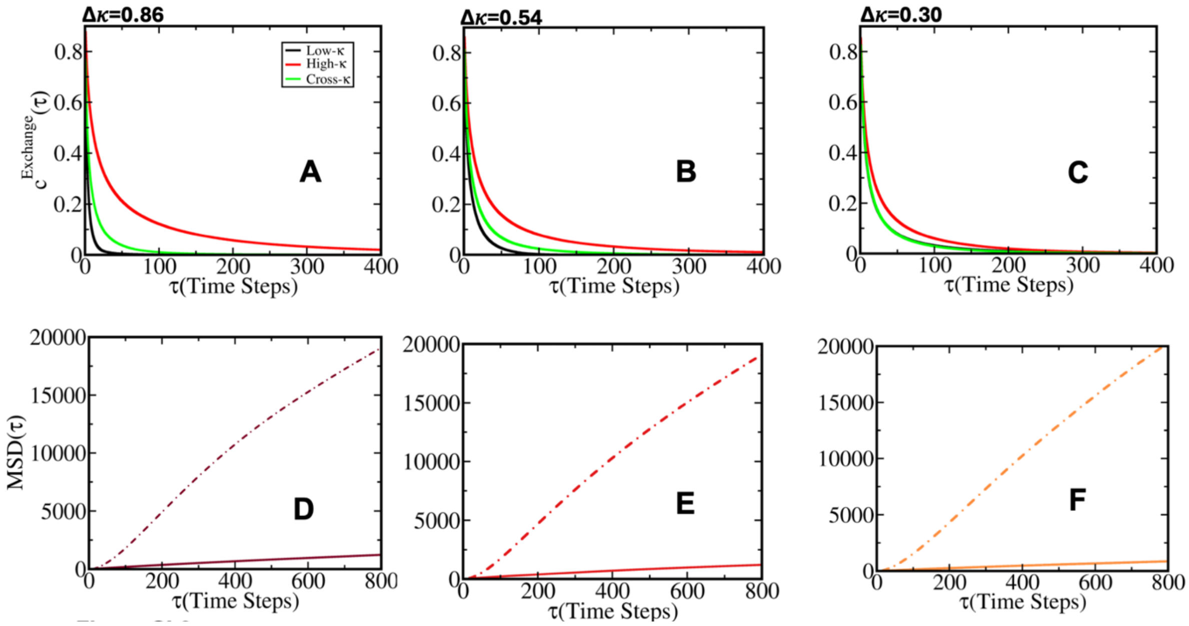

**Figure S6: Representative time correlation of contact exchange in droplet** phase among polymers at T/Tc=0.3 and $x_{Low-\kappa} = 0.5$ for sequence pairs (A) Δκ=0.86 (sequence κ=1.0 and 0.14) (B) Δκ=0.54 (sequence κ=0.77 and 0.23) (C) Δκ=0.30 (sequence κ=0.65 and 0.35). Each panel consists variants of contact lifetime correlation namely among polymers of low-κ (black), high-κ (red) and cross interaction among high and low κ polymers (green). A comparative analysis of average mean squared displacement (MSD) of polymers in droplet phase and bulk at T/Tc=0.3 and $x_{Low-\kappa} = 0.5$ for differential sequence pair condensates (A) Δκ=0.86 (sequence κ=1.0 and 0.14) (B) Δκ=0.54 (sequence κ=0.77 and 0.23) (C) Δκ=0.30 (sequence κ=0.65 and 0.35) at T=0. and $x_{Low-\kappa} = 0.5$ for sequence pairs.

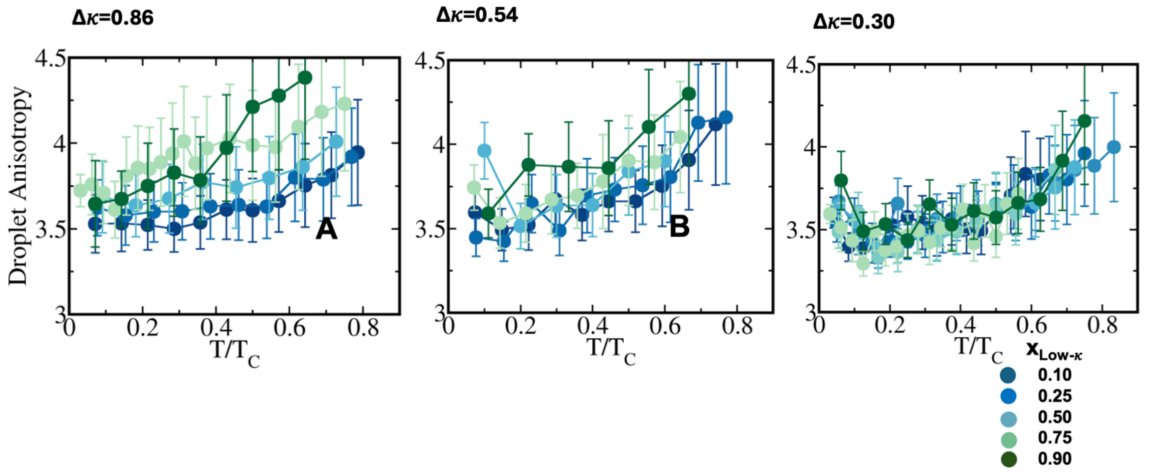


**Figure S7: Shape anisotropy of droplets composed by differential sequence pairs along temperature scaled to criticality (A)** Δκ=0.86 (sequence κ=1.0 and 0.14) (B) Δκ=0.54 (sequence κ=0.77 and 0.23) (C) Δκ=0.30 (sequence κ=0.65 and 0.35). Different mixing fractions has been shown in each panel ranging from $x_{Low-\kappa}$=0.1 to 0.9 with blue to green.
